# Databiology Lab CORONAHACK: Collection of Public COVID-19 Data

**DOI:** 10.1101/2020.10.22.328864

**Authors:** Juan Caballero Pérez, José Manuel Caballero Contreras, Carlos de Blas Pérez, Felipe Leza Alvarez

**Affiliations:** Databiology Ltd., Oxford, OX4 4GA, UK

**Keywords:** COVID-19, SARS-CoV-2, repository, data mining

## Abstract

COVID-19 has had an unprecedented global impact in health and economy affecting millions of persons world-wide. To support and enable a collaborative response from the global research communities, we created a data collection for different public sources for anonymized patient clinical data, imaging datasets, molecular data as nucleotide and protein sequences for the SARS-CoV-2 virus, reports of count of cases and deaths per city/country, and other economic indicators in Databiology Lab (https://www.lab.databiology.net/) where researchers could access these data assets and use the hundreds of available open source bioinformatic applications to analyze them. These data assets are regularly updated and was used in a successful virtual 3-day hackathon organized by Databiology Ltd and Mindstream-AI where hundreds of attendees to work collaboratively to analyze these data collections.

## Introduction

Coronavirus are a large RNA virus family, they are pathogens involved in human and animal diseases [1] and the recent emergence of a novel coronavirus in an outbreak in Wuhan, China has impacted almost all countries across the world as the called 2019-nCoV or COVID-19 pandemic. Initial sequencing analysis reported this new virus is a betacononavirus similar to the severe acute respiratory syndrome-related coronavirus (SARSr-CoV), therefore it was named as SARS-CoV-2 [2].

The COVID-19 disease could drastically affect or not the infected patient, there are many factors which can enhance the symptoms and co-morbidities such as cardiovascular, cerebrovascular, diabetes, obesity, and other diseases [3].

Since December 2019, multiple cases of COVID-19 have been reported, initially in the origin center in Wuhan in China’s Hubei Province, with a fast spread across the country and foreign countries. COVID-19 symptoms are diverse and infected patients could manifest fever, fatigue, dry cough, myalgia, diarrhea, and anosmia [4]. Laboratory tests can confirm the disease by viral nucleic acid detection [5].

Multiple treatments and vaccines are being developed to reduce the mortality and impact of the COVID-19 disease; however, this pandemic had already impacted humanity and legions of researchers are continuously looking into new insights to help fighting this disease. In aims to facilitate research, multiple data sets has been shared from diverse organizations and individuals, making it hard and difficult to have a good overview of the available and requiring multiple skills to retrieve, store, manage, clean-up, normalize and analyze those data sets.

Databiology Lab [https://www.lab.databiology.net/] is a research platform where researchers can register for free and access a curated list of projects loaded with COVID-19 data sets, including molecular data, cases and deaths per country and city, X-ray imaging data, 3D models for proteins, and more. In addition to the curated data the Databiology Lab platform also provides access to a large list of curated and commonly used tools in Bioinformatics as applications (Blast, Clustal Omega, Muscle, BWA, GATK, etc), interactive applications (MEGAN, UGENE, Emboss, ImageJ, IGV, etc), development frameworks (Jupyter, RStudio, Shelly) or developers can create and deploy their own tools using Databiology CIAO [https://learn.ciao.tools/]

In aims to showcase the data available in the platform, we perform a quick phylogenetic tree showing the genetic relationship between 100 SARS-CoV-2 proteins as encoded in the nucleic sequence isolated from patients.

## Methods

### Data integration

COVID-19 data was curated by searching in public sources, after initial screening to verify data availability, restrictions, format and structure, the sources were imported into Databiology Lab under 7 public projects, keeping the Databiology metadata structure (Subject → Episode → Sample → Extract → Resource) and resource original location whenever was possible.

### Phylogenic dataset

To build a Bayesian phylogeny, we used a selection of 100 amino acid sequences of SARS-CoV-2 from Project 407, the resources identifiers that we utilized was 6061100 – 6061199. As sequences are imported without any particular order, the selection can be considered a random subset of the total.

### Sequence alignment

We utilized the Muscle app (App ID: 20, https://www.lab.databiology.net/dbe/userlab/show-application.html?applicationId=20) to perform the multiple alignment for the sequences with the following parameters: neighbor-joining method for clustering and the maximum number of iterations was 2.

### Phylogenetic tree

To generate a Bayesian phylogenetic tree, we used MrBayes app (App ID: 214, https://www.lab.databiology.net/dbe/userlab/show-application.html?applicationId=214), with the following parameters: lset nst=1, rates=invgamma, mcmc ngen=10000, samplefreq=100, printfreq=100, diagnfreq=100, burnin=10, relburnin=YES, nchains=4, temp=0.5, stoprule=YES.

## Results

Databiology Lab was used to curate and index data and metadata in a Project (collaboration name space), while the original data remains stored in the original location, users can use such projects with different modes for access. The COVID-19 data sets have been set to be readable for all users in the Databiology Lab platform and the users can use their own Projects to run and share any analysis as they may decide. The following are the data sets made available in Databiology Lab:

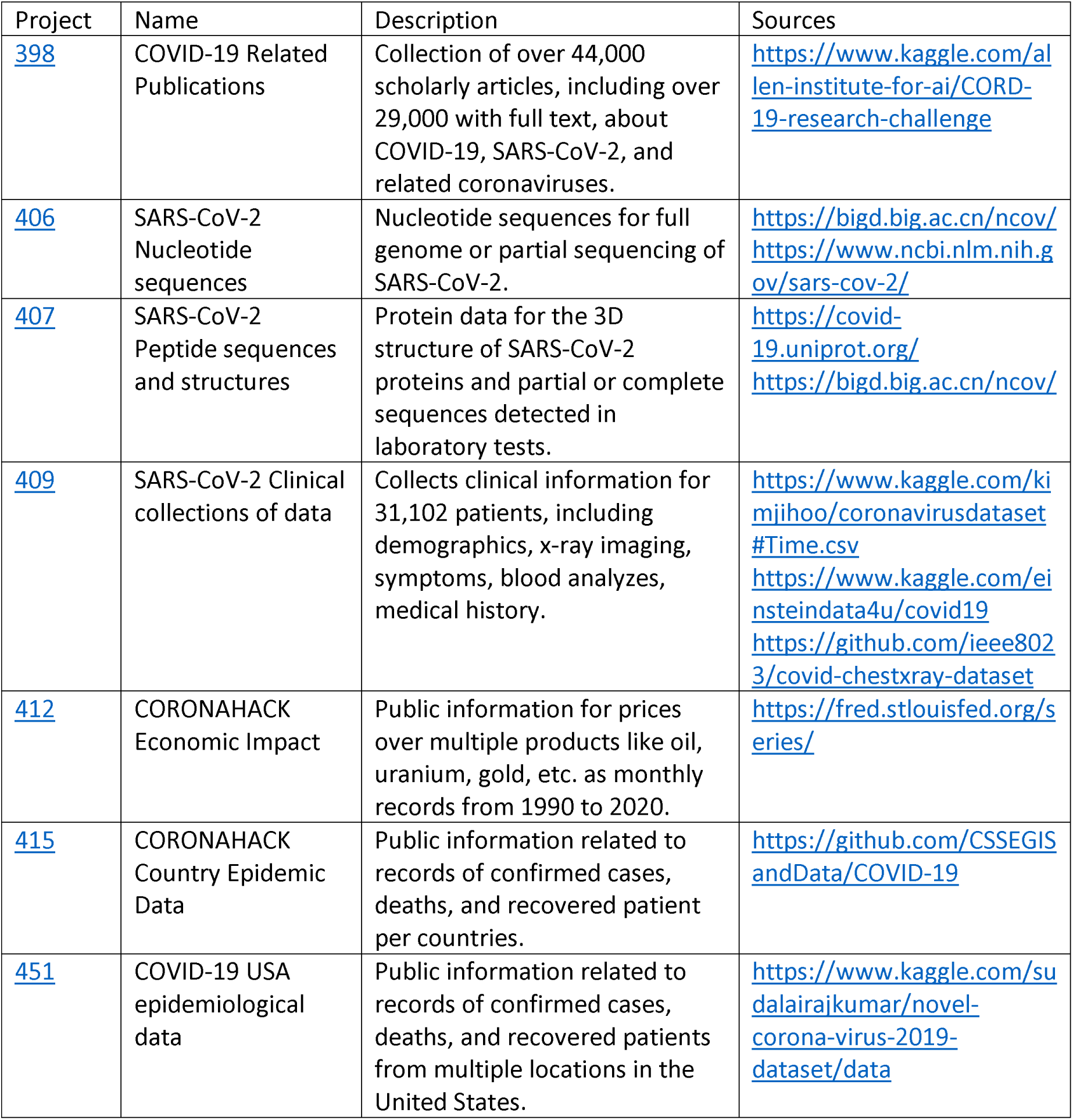

### Phylogenetic analysis

The analysis of the data in a phylogenetic tree of 100 sampled protein regions shows how multiple tools can be used to research using the provided data, Figure 1 shows the product of the analysis.

**Figure 1.**
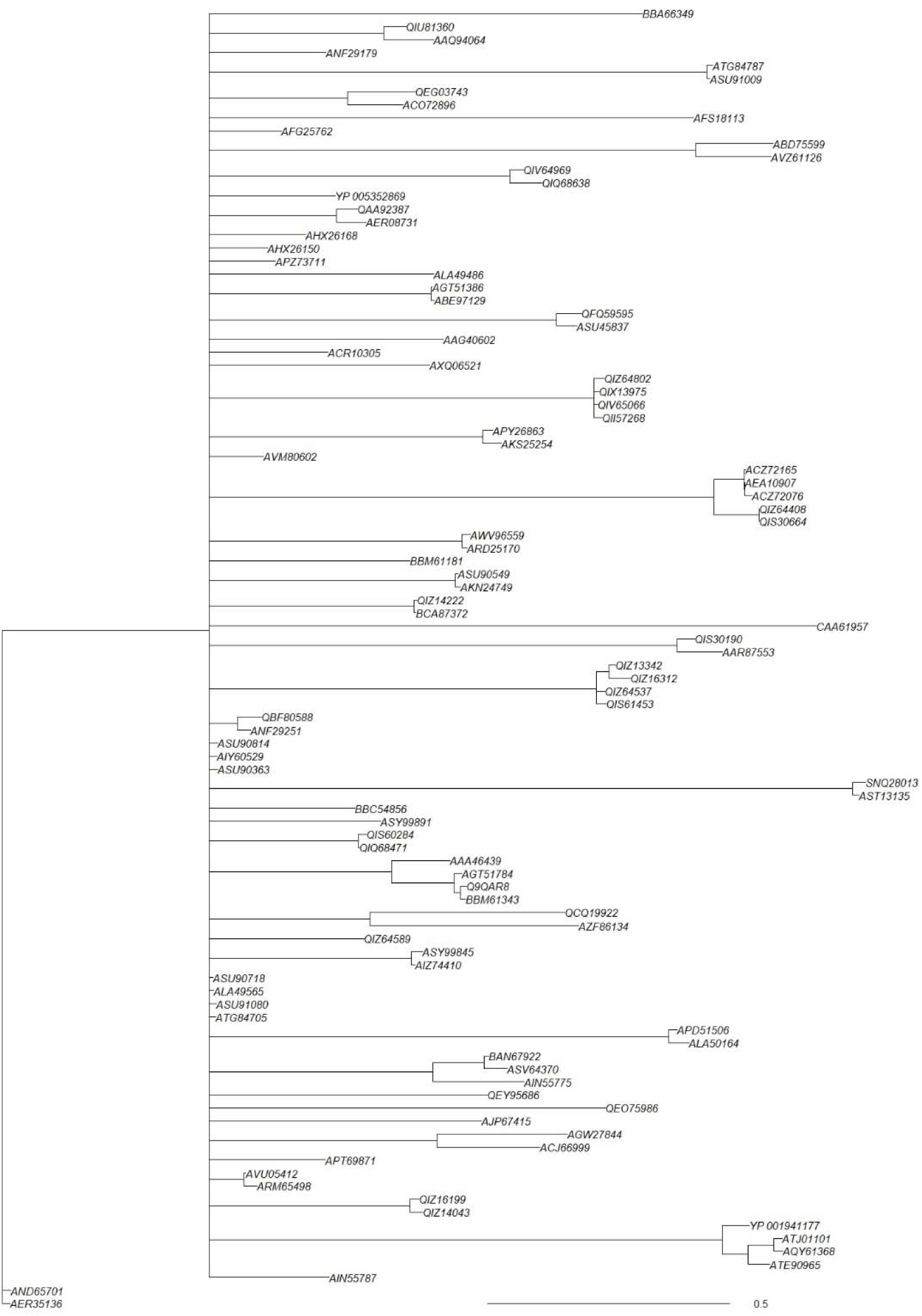
Phylogenetic tree for a subsampled 100 protein sequences of SARS-CoV-2 as detected in same number of nucleic samples obtained from COVID-19 patients.

## Discussion

COVID-19 impacted the world and after 10 months, data is still being generated and collected in order to better understand the disease and propose effective treatments. The data sharing and availability is required to facilitate research and encourage Open Science, using the FAIR principles (from findability, accessibility, interoperability, and reusability) [6] avoiding having partial and irreproducible results which cannot be evaluated or reproduced by other experts. Databiology products follow the FAIR principles, including data and analysis.

The available COVID-19 data sets in Databiology Lab are integrating multiple sources to reduce the time and skills needed to access the data enabling the researchers to focus on the analysis, as was demonstrated in the Hackathon [https://www.coronahack.co.uk/] where >350 Data Scientists and Researchers employed their skills to analysis the data provided, giving impressive results after few days of work [https://medium.com/@pauldowling/accelerating-scientific-collaboration-in-real-time-e1f682f54c87].

After the Hackathon, Databiology has continued to providing access to those data sets to any interested researcher without any cost and perform regular updates to ensure data is updated and new data sets are added as they become available.

We consider that making data available to other researchers is an opportunity to get bright minds to work in new creative ways to improve our understanding of COVID-19 and its impact in health, social, cultural and any other areas.

## Data availability

Data is available in Databiology Lab [https://www.lab.databiology.net/] in projects 398, 406, 407, 409, 412, 415, and 451.

## References

1. Chen Y., Liu Q., Guo D. Emerging coronaviruses: genome structure, replication, and pathogenesis. J Med Virol. 2020; 92:418–423

2. Mousavizadeh L., Ghasemi S. Genotype and phenotype of COVID-19: Their roles in pathogenesis. J Microbiol Immunol Infect. 2020; doi: 10.1016/j.jmii.2020.03.022.

3. Chen N., Zhou M., Dong X., Qu J., Gong F., Han Y. Epidemiological and clinical characteristics of 99 cases of 2019 novel coronavirus pneumonia in Wuhan, China: a descriptive study. Lancet. 2020; 395(10223):507–513

4. Li H., Liu Z., Ge J. Scientific research progress of COVID-19/SARS-CoV-2 in the first five months. J Cell Mol Med. 2020; 24(12):6558-6570. doi: 10.1111/jcmm.15364.

5. Huang C., Wang Y., Li X., Lofy K.H., Wiesman J., Bruce H. Clinical features of patients infected with 2019 novel coronavirus in Wuhan, China. Lancet. 2020; S0140–6736(20):30183–30185.

6. Wilkinson M.D., Dumontier M., Aalbersberg I.J., Appleton G., et al. The FAIR Guiding Principles for scientific data management and stewardship. Scientific Data 2016; 3:160018. doi:10.1038/sdata.2016.18

